# Amyloid-β precursor protein promotes tumor growth by establishing an immune-exclusive tumor microenvironment

**DOI:** 10.1101/2025.02.10.637339

**Authors:** Tao Yin, Guoping Wang, Zhehao Ma, Liuyang Wang, Rui Chen, Kun Xiang, Lianmei Tan, Yan Wang, Mengyang Chong, Yaosi Liang, Christopher C. Pan, Peter B. Alexander, Bryan Jian Wei Lim, Ergang Wang, Bushangqing Liu, Chengsong Yan, Qi-Jing Li, Xiao-Fan Wang

**Author notes:** These authors contributed equally (Tao Yin, Guoping Wang, Zhehao Ma). corresponding authors: Dr. Xiao-Fan Wang,; Dr. Qi-Jing Li.

## Abstract

During initiation and progression, cancerous tissue hijacks a series of elaborate tissue homeostatic mechanisms to avoid immune surveillance, including neuro-immune interactions. Here, we show that amyloid-β precursor protein (APP) and its β-cleavage product amyloid-β1-42 (Aβ1-42), well-known in the pathogenesis of Alzheimer’s disease (AD), are expressed in multiple cancer tissues. However, the oncogenic activity of APP is due to its E1 domain, instead of Aβ1-42. Mechanistically, APP restricts immune cells influx into tumor microenvironment (TME) and impairs CD8+ T cell and NK cell-based immunity, by dampening type I interferon (IFN) response in TME. We also provide proof-of-concept that vaccination targeting APP is effective for cancer prevention. Our current study reveals a previously unrecognized role of APP in cancer immune surveillance, and provides a new strategy for cancer prevention and treatment by targeting APP.

## Introduction

To maintain a physiological homeostasis, many tissue-policing mechanisms have been evolved to facilitate host defense^1,2^. Referred to as wounds that never heal, cancers exhibit disordered tissue homeostasis, and develop multiple strategies to evade host protective immune surveillance^3,4^. Especially noteworthy, optimal trafficking of T cells into tumor lesion represents a crucial step in the cancer-immunity cycle, which involves several sequential steps for complete tumor eradication^5^. Consistently, a T cell-inflamed TME is essential to generate a robust anti-tumor response and predicts better therapeutic response^6^. However, in general, this recruitment process is frequently impaired in the TME, which renders the failure of tumor growth control^7^. Poor intratumoral infiltration is also the barrier for effective immunotherapy in solid tumors, including immune checkpoint inhibitors^8^ and chimeric antigen receptor (CAR)-T cell therapy^9^. On the other hand, therapeutically restoring T cell recruitment could boost anti-tumor immune response by multiple approaches, such as adoptive cellular therapy^10^ and cancer vaccine^11^. Meanwhile, the determinants associated with the low frequency of tumor-infiltrating immune cells remain poorly understood. Given the importance of immune cell infiltration into TME, deciphering its mechanistic modulation is necessary.

In recent years, nervous system has emerged as one important modulator in TME^12^. A variety of tumors have been demonstrated to be innervated^13^, which could provide proliferative signaling^14^, support angiogenesis^15^, and induce T cell exhaustion^16^. To drive cancer progression, beyond directly interacting with nerve fibers, cancerous tissues frequently release diverse neuronal-related factors^17,18^, which hold essential functions for the development and activity of the nervous system^19^. Functionally, these tumor cell-intrinsic factors could be deployed to establish complex patterns of crosstalk between tumor cells and the surrounding immune cells. For example, we recently found that melanoma cells can hijack the immune privilege of nervous system. By secreting nerve growth factor (NGF), melanoma maintains an immune-excluded TME and suppresses the function of low-affinity CD8+ T cells; blockade of NGF signaling significantly sensitizes immunotherapy^20^. Therefore, targeting neural-tumor communication in the TME represents a novel approach for cancer therapy.

## Results

### APP and its β-cleavage pathway in human lung tumor tissues

APP is known to participate in the pathogenesis of AD as the source of the Aβ fragment^21^. As a single-pass transmembrane protein, APP presents a large extracellular domain followed by the juxta membrane sequence of Aβ, which further extends into the transmembrane domain. The cleavage of APP by either α-or β-secretase liberates its large ectodomain, which are respectively termed sAPPα and sAPPβ. The membrane-bound C-terminal stubs (C83 and C99) can be further processed by γ-secretase to generate the APP intracellular domain (AICD) and extracellular p3 or Aβ peptides (Fig 1A)^22^. In physiological conditions, APP cleavage mainly follows a non-amyloidogenic pathway; but it can proceed to an amyloidogenic β-cleavage in a pathological scenario, such as hypoxia^23^, which is a common feature of malignant tumors^24^. However, up to date, the expression of β-cleavage products was rarely documented in malignancy. To explore this possibility, we performed immunohistochemical (IHC) staining in cancer tissues for APP, Aβ1-42 and β-site APP cleaving enzyme 1 (BACE1), which initiates the amyloidogenic APP-processing pathway^25^. In both lung squamous carcinoma (LUSC) and adenocarcinoma (LUAD), tumor cells displayed a strong immunoreactivity to APP, while only a minority of stroma cells were APP-positive (Fig 1B, C, Supplementary Fig 1A). Interestingly, expression of Aβ1-42 and BACE1 was also observed in lung cancer tissues (Fig 1B, C), which was confirmed by antibodies against Aβ1-42 from different commercial sources (Supplementary Fig 1B, C). Furthermore, we observed a wide range of Aβ1-42 expression in multiple tumor tissues, including lung, breast and colon cancers (Supplementary Fig 1D-H), suggesting the presence of activated amyloidogenic APP-processing pathway in a broad spectrum of malignant conditions.

**Fig. 1.**
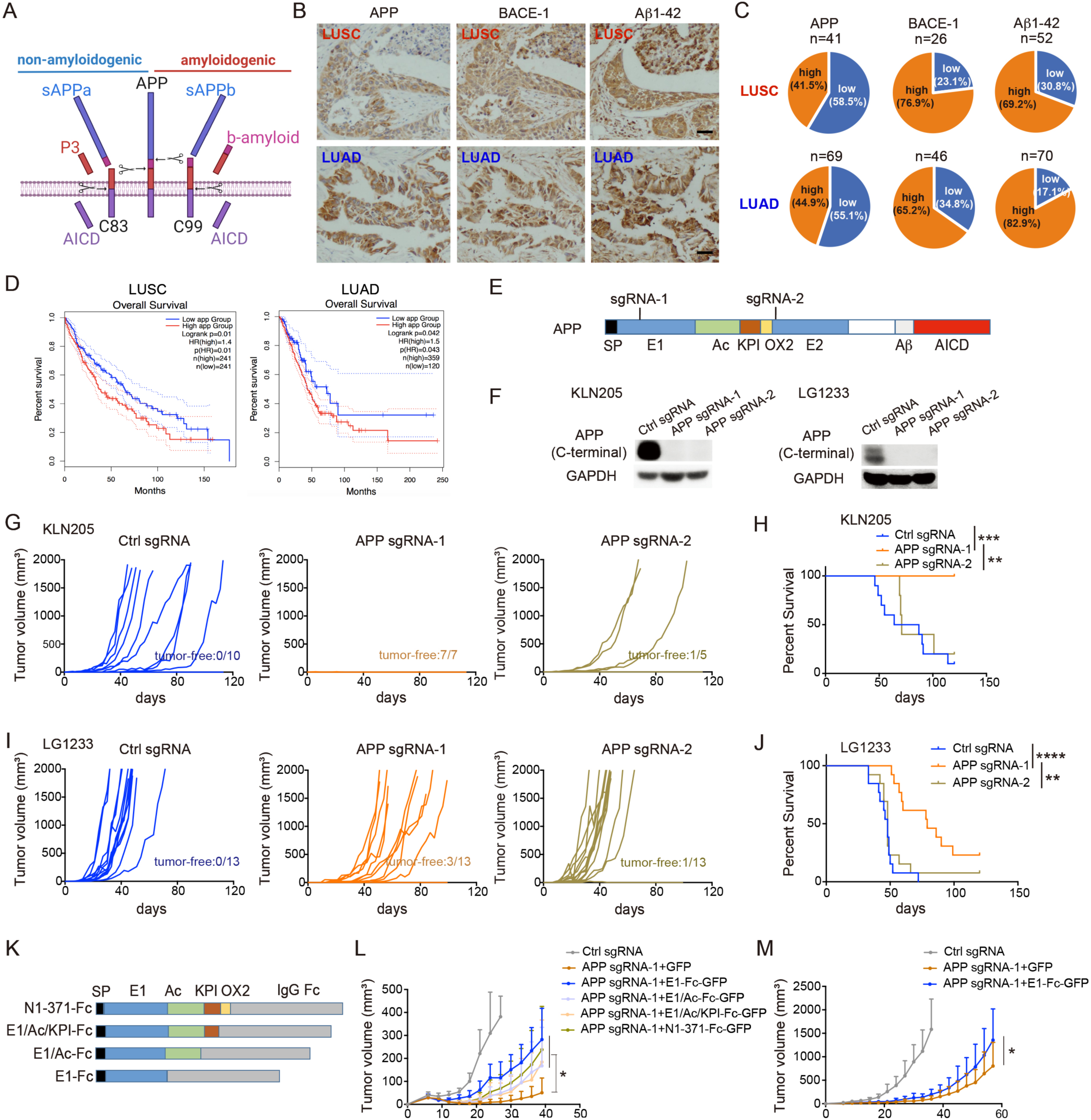
APP’s E1 domain promotes tumor growth. A, Schematic representation of the canonical APP processing pathways. B, Immunohistochemical staining of APP, BACE-1 and Aβ1-42 in human LUSC and LUAD. Scale bars, 50 μm. C, Quantification of APP (n=41 for LUSC, n=69 for LUAD), BACE-1 (n=26 for LUSC, n=46 for LUAD) and Aβ1-42 (n=52 for LUSC, n=70 for LUAD) expression as in (B). D, Human lung cancer gene expression and associated clinical data were retrieved from the GEPIA2 portal. Red and blue curves represent the high and low expression of APP, respectively. E, Schematics of the APP targeting site by single-guide RNA. F, Western blot using anti-APP C-terminal antibody to detect the knockout efficiency of sgRNAs. G-H, KLN205 cells with control sgRNA, APP sgRNA-1 and APP sgRNA-2 were injected subcutaneously into DBA/2J mice. n=7-14. I-J, LG1233 cells with control sgRNA, APP sgRNA-1 and APP sgRNA-2 were injected subcutaneously into C57BL/6J mice. Tumor volume was measured and survival curve was plotted. n=13. K, Schematics of domain rescue strategy. L, KLN205 cells with control sgRNA, APP sgRNA-1 and APP sgRNA-1 plus indicated APP domain-Fc were injected subcutaneously into DBA/2J mice. Tumor volume was measured. n=7-8. M, LG1233 cells with control sgRNA, APP sgRNA-1 and APP sgRNA-1 plus APP E1 domain-Fc were injected subcutaneously into C57BL/6J mice. Data were combined from two independent experiments. n=12-14. Data are mean ± s.d. \**P* < 0.05, \*\**P* < 0.01, \*\*\**P* < 0.001, \*\*\*\**P* < 0.0001.

We next evaluated the expression of APP and BACE-1 in a panel of human lung cancer cell lines and found the majority of them express APP and BACE-1, as revealed by western blot analysis (Supplementary Fig 1I), which is in line with our IHC data (Fig 1B, C). Consistently, the expression of APP and BACE-1 proteins was also observed in murine lung cancer cells, including LUSC cell line KLN205 (derived from bronchial carcinogen exposure^26^), and LUAD cell line LG1233 (containing K-ras mutation and P53-loss derived from C57BL/6 mice^27^) (Supplementary Fig 1J). APP signal was also detected by the anti-KPI and anti-OX2 antibodies recognizing two motifs in its extracellular domain, suggestive of expression of the APP770 isoform in lung cancer cells (Supplementary Fig 1I-K).

Further, we analyzed multiple datasets from Cancer Genome Atlas (TCGA) using the GEPIA2 online portal^28^, where a higher level of APP expression is associated with worse prognosis in cancer patients with LUSC or LUAD (Fig 1D). Altogether, these data suggest that APP and its β-cleavage products may have an oncogenic function.

### APP’s E1 domain, instead of the Aβ domain, promotes tumor growth

To explore a potential role of APP in oncogenesis, we firstly evaluated the functional significance of Aβ1-42 expression in tumor growth. The BACE1 inhibitor verubecestat has been shown to reduce Aβ1-42 production in animal models and in patients with AD^29^. However, when KLN205 tumor-bearing DBA/2J mice and LG1233 tumor-bearing C57BL/6 mice were dosed daily with verubecestat, we did not observe any anti-tumor efficacy of this inhibitor (Supplementary Fig 2A, B). We next administered an Aβ1-42 vaccine, which has also been used to reduce Aβ1-42 in preclinical models and AD patients^30,31^. Similarly, this approach did not provide any noticeable anti-tumor efficacy (Supplementary Fig 2C, D). In line with this, as revealed by the IHC staining data, we noticed that Aβ1-42 had a minimal effect on patient prognosis in multiple human tumor tissues, although there was a weak correlation in LUSC (Supplementary Fig 2E-H). These data indicate that Aβ1-42 does not exert a significant biological effect on lung tumor growth, so that its elevation in tumors might represent a byproduct of APP’s metabolism.

Since APP itself is a poor prognostic indicator (Fig. 1D), we therefore set out to determine whether APP is functionally important for tumor growth. As a first approach, we downregulated APP using RNA interference (Supplementary Fig 2I, K). From the *in vitro* experiments, both KLN and LG1233 murine tumor cells grew slightly slower than their nontargeting control counterparts (Supplementary Fig 2J, L). This is in line with other reports showing that APP could promote certain cancer cell proliferation^32,33^. However, tumor growth was much more strongly reduced in immune-competent syngeneic mouse models (Supplementary Fig 2M, N), indicating that APP might have other functions in addition to stimulating cell growth.

As a second approach, to completely deplete APP in tumor cells, we generated APP knockout (KO) cell lines using CRISPR-Cas 9-mediated targeting of the *App* gene. Specifically, two non-overlapping guide RNAs (sgRNA) were developed targeting exon 4 (sgRNA-1) and exon 9 (sgRNA-2) (Fig 1E). Both sgRNAs introduced frameshift indels near expected cut-site (Supplementary Fig 2O, P) and eliminated the expression of full-length APP, as revealed by Western blot using an anti-C-terminal APP antibody (Fig 1F). Intriguingly, when the cells were injected subcutaneously into the syngeneic mice, a dramatic discrepancy was observed. While APP sgRNA-2 slightly reduced tumor growth, the anti-tumor effects were much more pronounced in mice engrafted with the APP sgRNA-1 tumor cells. Indeed, 100% of mice injected with 150K APP sgRNA-1 KLN205 cells failed to display measurable tumor growth after implantation into immune-competent DBA2/J mice (Fig 1G, H). Similarly, sgRNA-1 but not sgRNA-2 significantly slowed down LG1233 tumor growth in C57BL/6J mice, resulting in a substantial survival benefit (Fig 1I, J). These unexpected results hinted that there was a huge difference between sgRNA-1– and sgRNA-2-mediated editing of the *App* gene. Since we used the single-cut CRISPR technology, sgRNA-2, which targets exon 9, may create a functional APP protein fragment (there are 371 aa upstream from the target site), whereas sgRNA-1 targeting exon 4 may eliminate expression of the entire APP. To test this postulation, we further conducted rescue experiments in the same two tumor models.

The ectodomain of APP770 isoform contains several functional domains defined previously by studies of APP in the neuronal system^22^, and the N-terminal fragment upstream from the sgRNA-2 target site (N1-371) contains the E1 domain, Acidic (Ac) domain, Kunitz protease inhibitor (KPI) domain and OX2 domain (Fig 1E). To test the functionality of these domains, we truncated them from APP’s C-terminus and conjugated these pieces with the IgG Fc fragment to stabilize the chimeric products (Fig 1K, Supplementary Fig 2Q). All Fc-fused domains, including E1, E1/Ac, E1/Ac/KPI, and N1-371 (which contains E1/Ac/KPI/OX2 domains), partially but significantly rescued KLN205 APP sgRNA-1 (hereafter designated as APP KO) tumor growth in wide-type (WT) mice, indicating that the E1 domain of APP may have a critical role in promoting tumor growth (Fig. 1L). The E1 domain alone also effectively, albeit partially, rescued the growth of LG1233 APP KO tumors (Fig 1M, Supplementary Fig 2R). Together, these data suggest that APP’s E1 domain, instead of the Aβ1-42 domain, promotes tumor growth in syngeneic mouse models.

### Remolding of tumor microenvironment by APP inactivation

To determine the biological nature of altered tumor features by APP deprivation, we performed RNA sequencing (RNA-seq) analysis of bulk tumor tissues. In these two different tumor models, multiple important hallmark features, including angiogenesis, hypoxia, apoptosis and cell cycle, were not changed in parallel; however, enhanced expression of gene signature involved in inflammatory response was always observed after APP inactivation (Supplementary Fig 3A, B), indicating that host immune system is engaged in the retarded APP-deficient tumor growth. Consistently, when 150K control and APP KO KLN205 tumor cells (the same number of injected tumor cells as in WT mice, in Fig. 1G, H) were transplanted into T-cell-compromised athymic nude mice, APP KO tumor cells grew out, although at a slower rate than the control cells (Fig 2A), indicating that T cells play a dominant role in contributing to APP-deficiency induced tumor suppression. We also observed that APP KO LG1233 tumors grew much faster in TCRα-/-mice than that in WT mice (Fig 2B), pointing to a key role of T cells in APP’s pro-tumor activity. To illustrate the immune landscape changes induced by APP deprivation, we utilized LG1233 bulk tumor tissue RNA-seq to conduct immune deconvolution. This analysis showed a global enrichment of immune cells in APP KO tumors (Fig 2C), which was confirmed by immunofluorescence staining and flow cytometry: APP depletion fostered a massive infiltration of CD45+ leukocytes into the TME (Fig 2D, E, Supplementary Fig 3C-E). Specifically, intratumoral enrichment of lymphocytes, including CD4+ T cells, CD8+ T cells, NK cells, and myeloid cells was observed in APP KO tumors (Fig 2F-K, Supplementary Fig 3F). Thus, these data demonstrate that APP inactivation augments immune cells infiltration and promotes an immune “hot” TME. Furthermore, as revealed by bioinformatic analysis in lung cancer patients, *APP* expression has a negative correlation with presence of CD8+ T cells and NK cells, supporting the clinical relevance of our preclinical findings (Supplementary Fig 3G, H).

**Fig. 2.**
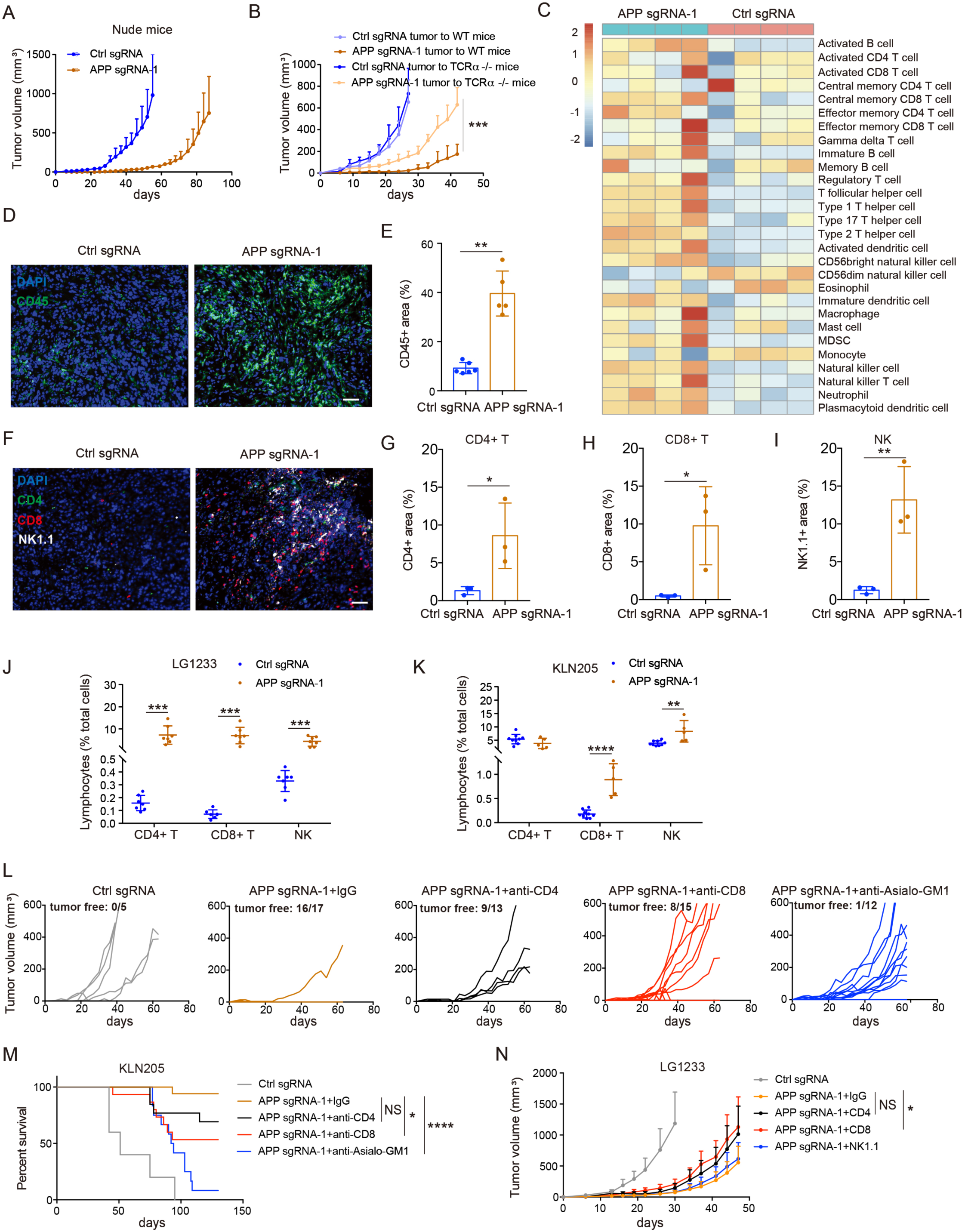
APP remolds tumor microenvironment. A, Tumor growth curves of athymic nude mice subcutaneously injected with KLN205 cells with control or APP sgRNA-1. n = 5. B, Tumor growth curves of C57BL/6J and TCRa-/-mice subcutaneously injected with LG1233 cells with control or APP sgRNA-1. n = 7-8. C, LG1233 cells with control or APP sgRNA-1 were injected into C57BL/6J mice. Heat map of immune deconvolution for Ctrl or APP sgRNA-1 LG1233 tumor tissues. n=4. D-E, LG1233 cells with control or APP sgRNA-1 were injected into C57BL/6J mice. CD45 immunofluorescence of Ctrl or APP sgRNA LG1233 tumor tissues. Nuclei were counterstained with DAPI. Scale bars, 50 µm. F-I, CD4, CD8 and NK1.1 immunofluorescence of Ctrl or APP sgRNA LG1233 tumor tissues. Nuclei were counterstained with DAPI. Scale bars, 50 µm. J, Control or APP sgRNA-1 LG1233 cells were subcutaneously injected into C57BL/6J mice. The percentage of lymphocytes in tumor microenvironment were analyzed by flow cytometry in Ctrl or APP sgRNA LG1233 tumor tissues. n=7. K, Control or APP sgRNA-1 KLN205 cells were subcutaneously injected into DBA/2J mice. The percentage of lymphocytes in TME were analyzed by flow cytometry in Ctrl or APP sgRNA LG1233 tumor tissues. n=9 for Ctrl, n=5 for KO. Each dot represents pooled sample from two small tumors. Data combined from two independent experiments. L-M, Tumor growth curves (L) and survival (M) of DBA/2J mice subcutaneously injected with Ctrl or APP sgRNA-1 KLN205 cells and treated with lymphocytes depletion antibodies. n=5-17. N, Tumor growth curves of C57BL/6J mice subcutaneously injected with Ctrl or APP sgRNA-1 LG1233 cells and treated with lymphocytes depletion antibodies. n=8-9. Data are mean ± s.d. \**P* < 0.05, \*\**P* < 0.01, \*\*\**P* < 0.001, \*\*\*\**P* < 0.0001.

Functionally, to determine whether increased effector lymphocytes are responsible for APP deprivation-mediated anti-tumor response, we individually depleted lymphocyte subsets (Supplementary Fig 4A-D). Depletion of CD8+ T and NK cells in immune-competent mice significantly diminished the APP KO-mediated protection against KLN tumors and significantly reduced the survival advantage provided by APP inactivation, whereas the effect of CD4+ T cell depletion on tumor growth was modest (Fig 2L, M). Similarly, the absence of CD8+ T cells in C57BL/6 mice also partially restored the growth of LG1233 APP KO tumors (Fig. 2N), demonstrating the accumulation of endogenous CD8+ T cells is essential for APP-deficient tumor control. In line with this, mice with their primary LG1233 APP KO tumors removed were resistant to subsequent re-challenge by inoculation of the parental tumor cell lines (Supplementary Fig 4E, F), but not unrelated B16 melanoma cells (Supplementary Fig 4G), pointing out the generation of a specific anti-tumor memory response after APP deprivation; this is also consistent with the increased memory CD8+ T cells in APP KO tumors (Fig 2C). These data suggest that APP disruption not only resulted in a robust anti-tumor response by establishing an immune “hot” TME, but also induced a durable and specific anti-tumor memory response.

### APP dampens tumor-cell intrinsic type I interferon response

Chemotaxis mediated by chemokines is critical for directing immune cell influx into the TME, thereby enabling effective antitumor response^34^. In particular, Cxcl9/10 are critical for intratumoral infiltration of effector cytotoxic cells^35,36^, and predominantly associated with anti-tumorigenic immune response^34^. To gain further insight into how APP impacts the immune cell network within TME, we examined the expression profiles of important chemokines. Through RNA-seq analysis, we found that there was a significant increase in the expression of Cxcl9/10 following APP ablation (Fig 3A). Functionally, we observed that neutralizing antibody against Cxcr3, a shared Th1-associated chemokine receptor by CXCL9/10/11^37,38^, significantly abolished the elevated immune cell chemotactic activities induced by APP deprivation (Fig 3B). Consistently, Cxcr3 blockade reversed APP-deficient tumor growth (Fig 3C, D). These data indicate that APP installs an immune “cold” TME by silencing the activity of Cxcl9/10-Cxcr3 chemotactic axis.

**Fig. 3.**
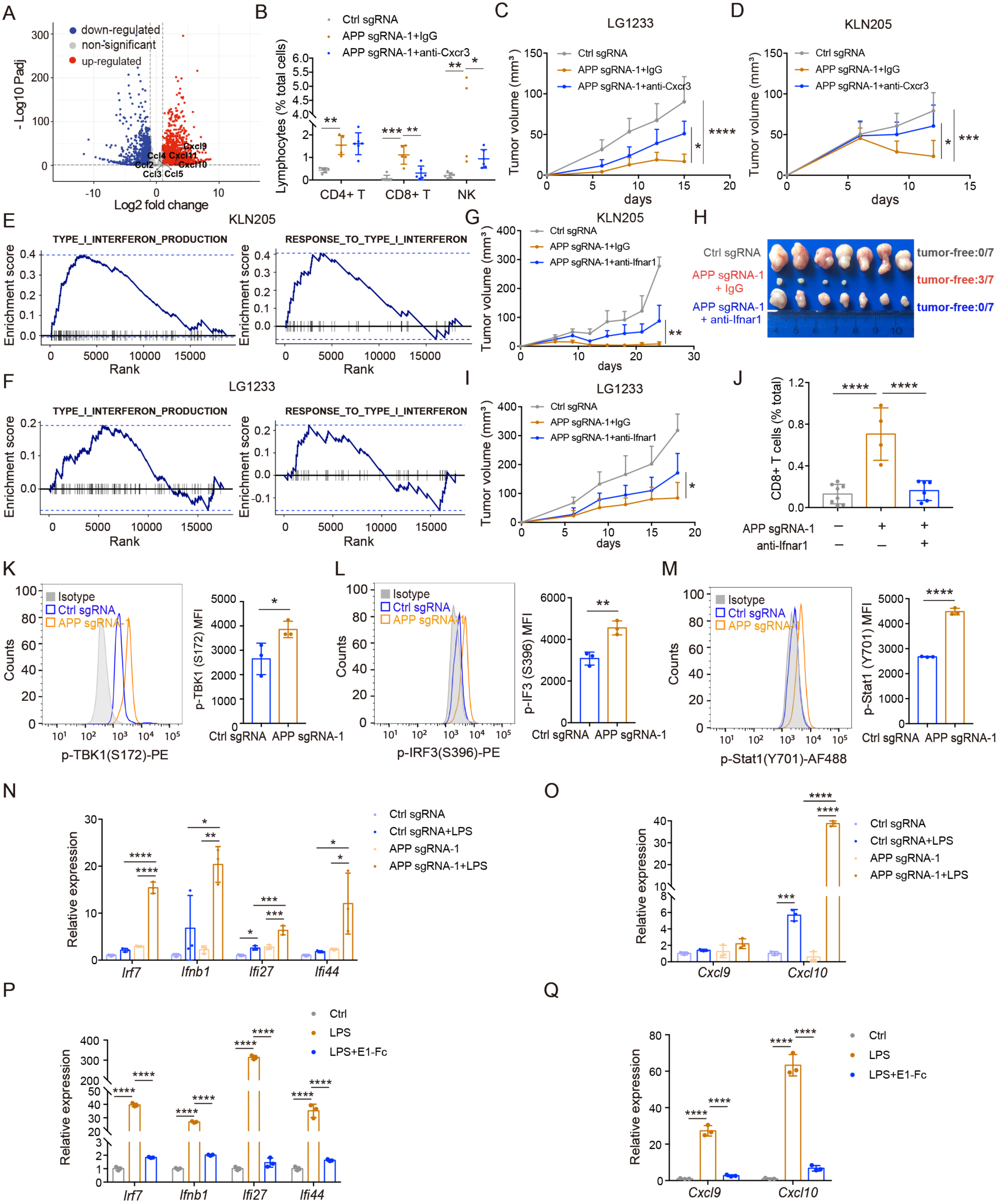
APP restrains type I interferon response. A, Volcano plot showing increased (red) and decreased (blue) chemokine gene expression (FDR-adjusted *P* < 0.05) in APP sgRNA-1 versus Ctrl sgRNA LG1233 tumors. The significantly differential chemokines were labelled. B, Ctrl or APP sgRNA-1 LG1233 tumor cells were injected into male C57BL/6J mice and treated with Cxcr3 neutralizing antibodies. The percentage of CD4+ T, CD8+ T and NK cells were analyzed by flow cytometry. n=6. C, Ctrl or APP sgRNA-1 LG1233 tumor cells were injected into C57BL/6J mice and treated with Cxcr3 neutralizing antibodies. Tumor growth curves were plotted. n=6. D, Tumor growth curves of DBA/2J mice subcutaneously injected with Ctrl or APP sgRNA-1 KLN205 tumor cells and treated with Cxcr3 neutralizing antibodies. n=6-8. E-F, Gene set enrichment analysis of RNA-seq data from bulk KLN205 (E) and LG1233 (F) tumor tissues. Type I interferon signaling was compared between APP sgRNA-1 versus Ctrl sgRNA tumors. G-H, Ctrl or APP sgRNA-1 KLN205 tumor cells were injected into male DBA/2J mice and treated with Ifnar1 neutralizing antibodies. Tumor growth curves (G) and representative photos of tumors (H). n =7. I, Tumor growth curves of C57BL/6J mice subcutaneously injected with LG1233 cells with control or APP sgRNA-1 and treated with Ifnar1 neutralizing antibodies. n = 7-9. J, Ctrl or APP sgRNA-1 KLN205 tumor cells were injected into male DBA/2J mice and treated with Ifnar1 neutralizing antibodies. The percentage of CD8+ T were analyzed by flow cytometry. K, the expression level of phosphorylated TBK1 (S172) in Ctrl sgRNA and APP sgRNA-1 LG1233 tumor cells was evaluated by flow cytometry. L, The expression level of phosphorylated IRF3 (S396) in Ctrl sgRNA and APP sgRNA-1 LG1233 tumor cells was evaluated by flow cytometry. M, The expression level of phosphorylated Stat1 (Y701) in Ctrl sgRNA and APP sgRNA-1 LG1233 tumor cells was evaluated by flow cytometry. N-O, Ctrl or APP sgRNA-1 LG1233 tumor cells were treated with LPS for 2 hr. The expression of type I interferon (N) and chemokines (O) was assessed by RT-PCR. P-Q, WT LG1233 tumor cells were treated with recombinant APP E1-Fc peptide for 12 hr, followed by treatment with LPS for 2 hr. The expression of type I interferon (P) and chemokines (Q) was assessed by RT-PCR. Data are mean ± s.d. \**P* < 0.05, \*\**P* < 0.01, \*\*\**P* < 0.001, \*\*\*\**P* < 0.0001.

Considering Cxcl9/10 was IFN-driven chemokines^39^, we postulated that APP might regulate Cxcl9/10 by suppressing IFN signaling, which is necessary for effective T cell infiltration^40^. Indeed, in our RNA-seq data, gene set enrichment analysis (GSEA) revealed a marked enrichment of type I IFN (IFN-I) production and response gene signature in APP-deficient tumors (Fig 3E, F). We next explored whether APP KO-induced type I IFN signaling was responsible for the *in vivo* tumor growth retardation. IFN-Is signal through their common cognate receptor, the interferon α/β receptor (IFNAR) which is composed of two subunits, IFNAR1 and IFNAR2^41^. We treated tumor-bearing mice with a neutralizing antibody against Ifnar1, ablation of which alone has been demonstrated to abrogate conventional IFN signaling^42^. Blockade of Ifnar1 markedly compromised the anti-neoplastic effects of APP inactivation (Fig 3G-I). Moreover, Ifnar1 blockade also reversed the influx of CD8+ T lymphocytes into APP KO tumor tissues (Fig 3J), consistent with the role of IFN-I in T cell infiltration in the TME^40^. These data suggest that APP suppresses anti-tumor immunity by impairing IFN-I signaling and its downstream chemokine expression.

As tumor cells are the main source of APP in tumor tissues (Fig 1B, Supplementary Fig 1A), we next probed whether tumor cell-derived APP could restrain the epithelial cell-intrinsic IFN-I signaling. Indeed, the phosphorylated TBK1 and IRF3, which are important for IFN-I production, was increased after APP deprivation (Fig 3K, L, Supplementary Fig 5A). In addition, elevated phosphorylation of Stat1confirmed the activation of IFN signaling pathways in APP KO tumor cells (Fig 3M). Moreover, when exposed to the danger signal represented by a TLR4 agonist lipopolysaccharide (LPS), APP KO tumor cells were more prone to produce IFN-I and Cxcl9/10 (Fig 3N, O). These data indicate that APP could desensitize tumor cells to IFN-I inducer and restrict tumor-intrinsic IFN-I production. We further tested whether APP’s E1 domain could suppress the IFN-I response by culturing tumor cells at low density to reduce cell-cell contact but provide exogenous recombinant APP E1-Fc peptide. In WT mouse and human tumor cells, pretreatment with recombinant APP E1-Fc, instead of the Fc fragment, significantly impaired the IFN-I response by TLR activation (Fig 3P, Q, Supplementary Fig 5B, C). Similarly, ectopic APP E1-Fc overexpression also inhibited the IFN-I response in WT tumor cells (Supplementary Fig 5D, E). This suppression was also observed in a panel of human cell lines (Supplementary Fig 5F, G). APP is known to present not only as a membrane protein, but also in soluble forms, namely sAPPα and sAPPβ, each of which contains the E1 domain^22^. Notably, both sAPPα and sAPPβ suppressed IFN-I response in tumor cells (Supplementary Fig 5H). Collectively, these data suggest that tumor cell-derived APP maintains an immune-exclusive TME through suppressing IFN-I response and its downstream chemokine expression.

### GSK3β mediates the suppressive effect of APP on IFN-I response

The magnitude and qualitative nature of IFN-I signaling are fine-tuned by opposing augmenting and suppressive factors^43^. To investigate the possible molecular mechanisms by which APP suppresses the IFN-I response, we compared the expression of important regulators of IFN-I signaling through analysis of bulk RNA-seq data. Among up-regulated modulators (Fig. 4A), glycogen synthase kinase 3β (GSK3β) is a multifunctional Ser/Thr kinase found in eukaryotes. Beyond involving in various essential cellular functions, such as glycogen metabolism, cell cycle regulation and cellular differentiation, GSK3β has been shown to participate in inflammation and immune response^44,45^. More importantly, GSK3β could promote TBK1 self-association and autophosphorylation at Ser172, which is critical for virus-induced IRF3 activation and IFN-β induction^46,47^. However, it is not clear whether GSK3β also positively regulates IFN-I response in malignant tumors. To address this question, we overexpressed constitutively active Gsk3β S9A into tumor cells (Supplementary Fig 6A, B), resulting in an enhanced IFN-I production and response *in vitro* (Supplementary Fig 6C, D). We next transplanted tumor cells overexpressing Gsk3β S9A into syngeneic mice and resultant tumor tissues were submitted for RNA-seq. By conducting gene ontology (GO) pathway analysis, we found that “response to interferon-beta” was the top enriched pathway in Gsk3β S9A-overexpressed tumors (Supplementary Fig 6E-G), Consistently, tumor growth was suppressed by Gsk3β S9A expression (Supplementary Fig 6H). These data suggest that Gsk3β favors IFN-I response and resultant anti-tumor response.

**Fig. 4.**
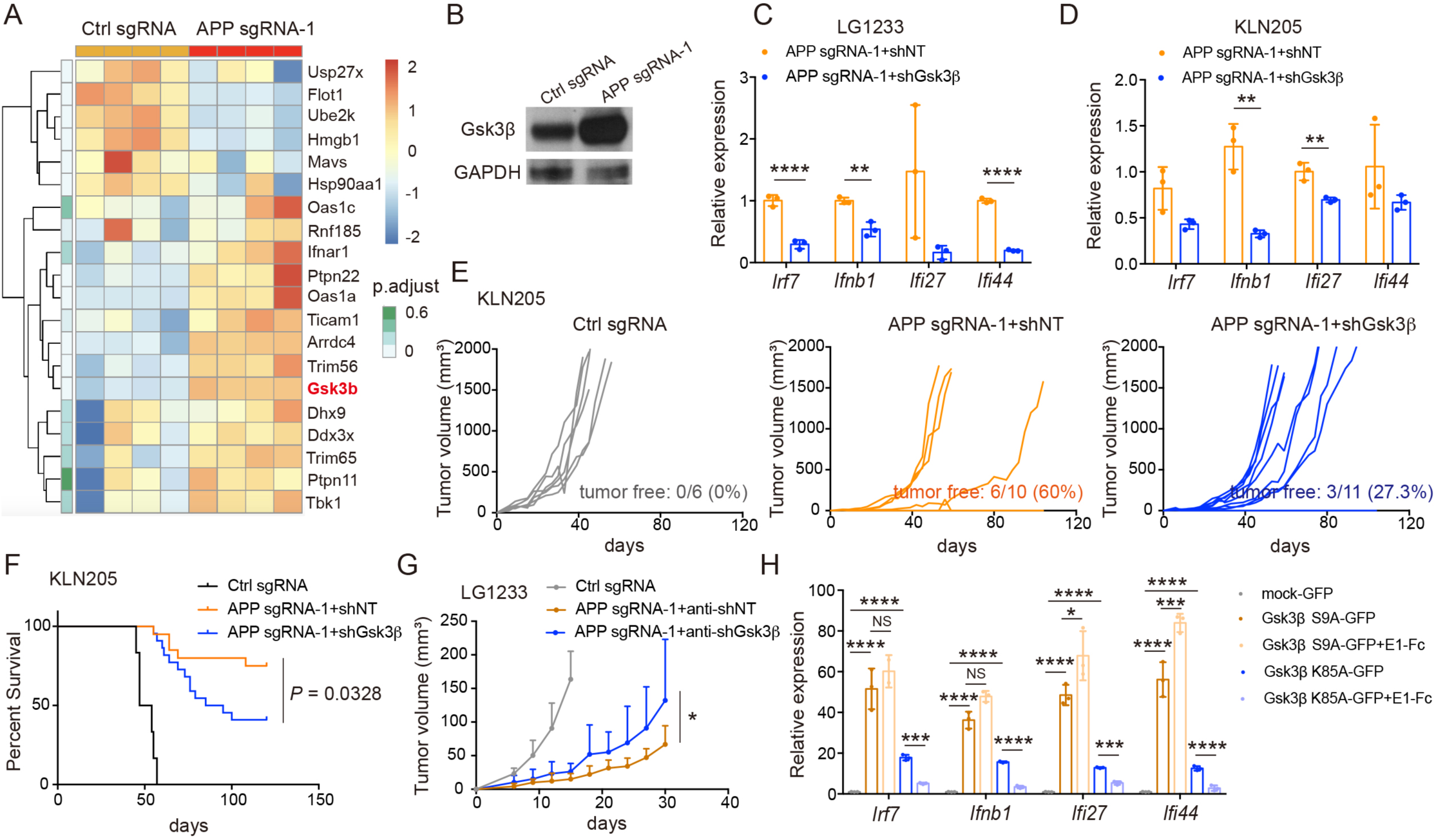
GSK3β mediated APP’s suppressive role in type I interferon response. A, Heatmap of IFN-I regulators in Ctrl or APP sgRNA-1 LG1233 tumor tissues. n=4. B, The expression of Gsk3β in APP sgRNA-1 LG1233 tumor cells assessed by western blot. C-D, LG1233 (C) and KLN205 (D) APP sgRNA-1 tumor cells with shGSK3β were treated with LPS, respectively, for 2 hr. The expression of IFN-I and related genes was evaluated by RT-PCR. E, The growth curve of APP sgRNA-1 plus shGsk3β KLN205 tumors. One representative data of two independent experiments. n=6-11. F, Survival curve of APP sgRNA-1 plus shGsk3β KLN205 tumors. Data were combined from two independent experiment. G, The growth curve of APP sgRNA-1 plus shGsk3β LG1233 tumors. H, LG1233 tumor cells with mock-GFP, Gsk3β S9A-GFP, Gsk3β K85A-GFP overexpression were treated with recombinant APP E1-Fc peptide for 12 hr, followed by treatment with LPS for 2 hr. The expression of IFN-I was assessed by RT-PCR. Data are mean ± s.d. \*\**P* < 0.01, \*\*\**P* < 0.001, \*\*\*\**P* < 0.0001.

Higher expression level of GSK3β was observed in APP KO tumor cells compared with WT cells (Fig. 4B). Moreover, GSK3β knock down by shRNA reduced IFN-I production (Fig. 4C, D, Supplementary Fig 6I, J) and partially restored the growth of APP-deficient tumor cells (Fig. 4E-G). Importantly, APP E1 treatment no longer suppressed IFN-I production in WT tumor cells with Gsk3β S9A overexpression (Fig 4H, Supplementary Fig 6K, L). However, APP E1 treatment still impaired IFN-I response in WT tumor cells with Gsk3β K85A ectopic expression (Fig 4H, Supplementary Fig 6M), which has an inactivated kinase activity but still can be degraded^46^, indicating that APP E1 regulated IFN-I signaling through GSK3β but independent of its kinase activity. This is consistent with another report that the effect of Gsk3β on virus-triggered induction of IFN-I is independent of its kinase activity^46^.

### APP’s E1 vaccine is effective for cancer prevention

In light of APP’s function in maintaining an immune “cold” TME, we explored whether blockade of APP might be potentially used for cancer prevention. Mindful that the E1 domain is located extracellularly, we focused on this domain as a targe for cancer vaccine, which provides the advantage of both ease of administration and control for individuals. To induce a stronger humoral response, we chose aluminum as the adjuvant, which selectively stimulates a Th2 immune response^48^. Due to central tolerance toward self-molecules, developing an effective self-antigen-based tumor vaccine is a difficult challenge; we therefore mutated several amino acid of APP E1 as an immunogen. Indeed, vaccination with mutant APP E1 peptide demonstrated an encouraging response in KLN205-bearing DBA/2J mice. Three doses of vaccination provided a striking preventative effect; notably, the survival of 70% vaccinated mice was prolonged beyond 4 months (Fig 5A, B). APP’s E1 peptide vaccine also afforded immune protection and triggered 80% tumor eradication in LG1233 tumor model in C57BL/6 mice (Fig 5C, D). To further test the generalizability of our E1 peptide vaccine, we extended our observation to two other tumor models with different genetic background: CT26 colon carcinoma in BALB/c mice and Py8119 breast cancer in C57BL/6 mice. The preventative immunization also induced a significant protection in both models (Fig 5E, F), suggesting that the APP E1 vaccine is effective for cancer prevention, irrespective of tumor histology or genetic background.

**Fig. 5.**
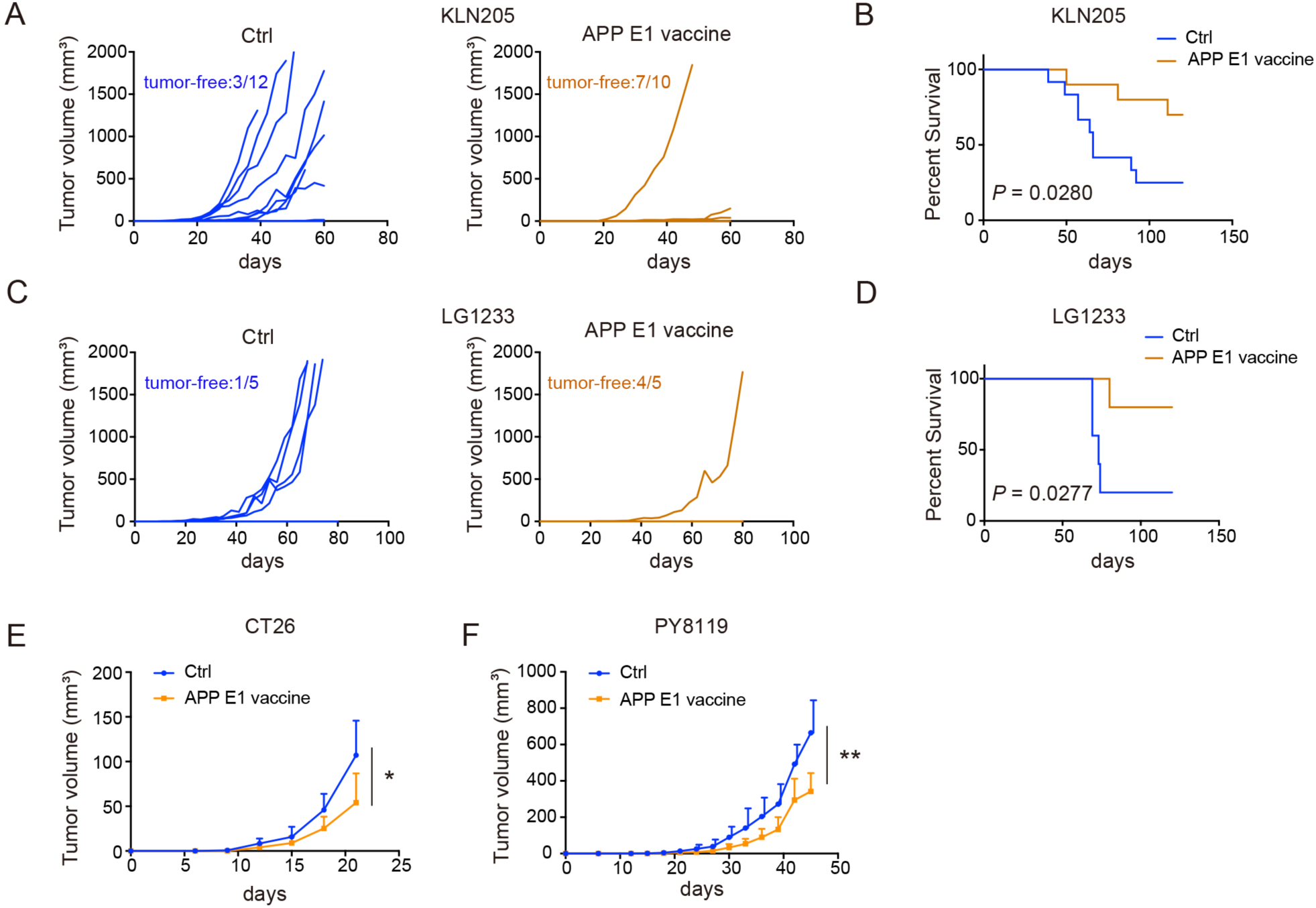
Targeting APP is effective for cancer prevention. A-B, DBA/2J mice were prophylactically vaccinated with mutant APP E1-Fc peptide along with adjuvant alum once a week for 3 weeks. Mice were then challenged with KLN205 cells. The growth kinetics of the individual tumors (A) and survival curve (B) of mice receiving vaccination. n=10-12. C-D, C57BL/6 mice were prophylactically vaccinated with mutant APP E1-Fc peptide along with adjuvant alum once a week for 3 weeks. Mice were then challenged with LG1233 cells. The growth kinetics (C) and survival curve (D) of LG1233 tumors. n=5. E, BALB/c mice were prophylactically vaccinated with mutant APP E1-Fc peptide along with adjuvant alum once a week for 3 weeks. Mice were then challenged with CT26 colon cancer cells. The tumor growth curves of mice receiving vaccination. n=6. F, C57BL/6 mice were prophylactically vaccinated with mutant APP E1-Fc peptide along with adjuvant alum once a week for 3 weeks. Mice were then challenged with PY8119 breast cancer cells. The tumor growth curves of mice receiving vaccination. n=7-9. Data are mean ± s.d. \**P* < 0.05, \*\**P* < 0.01.

## Discussion

Currently, the contribution of Aβ to the development of AD has been well established^21^. However, although much intense efforts have been dedicated to studies of amyloidogenic properties of the Aβ peptide and its targeted therapy, the therapeutic effect remains ineffective in slowing down the disease^49,22^, indicating other factors might be involved in the pathogenesis of AD. In the last decades, APP itself has been documented as a multifunctional protein implicated in several physiologic processes, including neuronal excitability, neurite outgrowth, synaptogenesis, synaptic plasticity and neuron survival^50^. Notably, as an evolutionarily conserved protein, APP actively participates in damage response in the central nervous system: APP is upregulated during brain injury^51,52^ and exerts neuroprotective effects following injury^53,54^. Outside the central nervous system, APP has been documented to be highly expressed in multiple cancer tissues and hinted to be involved in regulating tumor cell proliferation, differentiation and migration^55,56,32,57^. In this study, we have now provided more systemic evidence for the expression profiles of APP by malignant cells in comparison to stroma cells that could be the basis for tumor-intrinsic APP to exert a “protective” effect on cancer cells during coevolving with the host.

Furthermore, we found that targeting Aβ is not effective for cancer therapy, and there is a trivial prognostic value of Aβ expression in various cohorts of cancer patients, indicating that Aβ might be a byproduct of APP metabolism in the cancer context. Instead, APP’s E1 domain stands out as a critical player in promoting cancer growth via shaping the immunological landscape of the TME. By eliciting IFN-I response, APP inactivation is capable of altering T-cell trafficking and promoting an inflammatory TME in a CXCL9/10-dependent fashion. Coincidentally, APP has been shown to modulate inflammatory response in the central nervous system. For example, in Niemann-Pick disease type C, a neurodegenerative disease, loss of APP exacerbates the pathologic inflammation, even though the detailed molecular mechanism is unknown^58^. From this perspective, tumor cells may hijack APP’s physiological function in the brain to evade immune attack within the TME. Importantly, our findings reveal a novel function of APP in remolding TME, and highlight APP as a new appealing target for cancer immunotherapy. Indeed, as a proof-of-concept, vaccination against the E1 domain of APP could significantly prevent cancer growth. Lastly, given the contribution of immune response to AD^59^, and the uncovered novel function of APP in immunosurveillance by our study, the role of APP’s E1 domain in the pathogenesis of AD merits further study, which may potentially promote novel drug development for AD patients.

## Acknowledgement

We thank staffs at Duke University Flow Cytometry Shared Resource and Light Microscopy Core Facility for help with data acquisition. This work was supported by CA249726 from the NIH to X.-F.W.

## Methods

### Animals

All mouse experiments were performed with the approval from the Duke University Animal Care and Use Committee. All mice were housed under pathogen-free conditions at 72 °F, with a humidity of between 30 and 70%, a light/dark cycle of 12 h and *ad libitum* access to food and water. C57BL/6 mice, DBA/2J mice and TCRα-/-mice were bred in house. Athymic nude mice were obtained from Duke Breeding Core Facility.

### Cell culture

Human lung cell lines including A549, PC-9, HCC827, H1650, H2170 and H520 cells, were maintained in RPMI 1640 containing 10% FBS and 1% penicillin–streptomycin (GIBCO). Murine lung cancer LG1233 cell line was maintained in Dulbecco’s modified Eagle’s medium (DMEM) containing 10% FBS and 1% penicillin–streptomycin. Murine lung cancer KLN205 cell line was maintained in Minimum Essential Medium Eagle (Sigma) containing 10% FBS and 1% penicillin–streptomycin. HEK293FT cells were maintained in DMEM supplemented with 10% FBS and 1% penicillin–streptomycin. CT26 cells and B16F10 cells were maintained in RPMI 1640 containing 10% FBS and 1% penicillin–streptomycin (GIBCO). PY8119 were maintained in Ham’s F-12K media supplemented with 5% FBS and 1% penicillin–streptomycin.

### Generation of stable cell lines

CRISPR-Cas 9 technology was used to generate APP KO cell lines. Target sequences against mouse App (sgRNA 1: 5’-GCACTTGTCGGGCACGAGAA-3’; sgRNA 2: 5’-GTACTTGTCGACGGCGTCGG-3’) were designed in the Broad Institute GPP Web Portal. These sgRNA guides were synthetized as pairs of annealed oligos with overhangs cohesive to the digestion product of BmsBI restriction enzyme. Oligoduplexes were ligated to the lentiCRISPR v2 vector and transformed to DH5a bacteria. Vectors containing the sgRNA guides were transfected with pVSVG and pPAX2 to HEK293FT cells in order to produce lentiviral vectors. Lentiviral particles were produced by the transient transfection of HEK293FT cells using Lipofectamine™ 2000 (Thermo Fisher Scientific). Supernatant containing recombinant lentiviral particles were collected at 48 and 72 h after transfection. Tumor cells were cultured with supernatant containing viral particles followed by 2 μg/ml puromycin selection for 5 to 7 days. Single cell clones were generated and characterized. To confirm CRISPR-Cas 9 indel events near expected cut-site, DNA fragments were amplified from tumor cell populations with primers that anneal in introns flanking the targeted exons. PCR products were sequenced. To generate App or Gsk3b knockdown stable cell lines, App or Gsk3b shRNA lentiviral plasmid was purchased from Sigma-Aldrich or Genecopiea company, respectively. To prepare viral constructs, the vectors were co-transfected with pVSVG and psPAX2 plasmid. Lentiviral particles were produced by the transient transfection of HEK293FT cells using Lipofectamine™ 2000. Supernatant containing recombinant lentiviral particles were collected at 48 and 72 h after infection. Tumor cells were infected with supernatant containing viral particles followed by 2 μg/ml puromycin selection for 7 days. To generate stable cell lines with App domain overexpression, *App* gene with truncated domain was cloned into a MSCV retroviral vector. To generate stable cell lines with wide-type and mutant Gsk3b overexpression, *Gsk3b*, *Gsk3b S9A* and *Gsk3b K85A* gene was cloned into a MSCV retroviral vector. To prepare viral constructs, these vectors were co-transfected with pVSVG and pGag/Pol plasmid. Retroviral particles were produced by the transient transfection of HEK293FT cells using Lipofectamine™ 2000. Recombinant lentiviral particles were collected at 48 and 72 h after infection. Tumor cells were cultured with supernatant containing viral particles and GFP-positive cells were sorted out by flow cytometry.

### Murine tumor models

For *in vivo* experiments, a total of 2 × 10^3^, 1 × 10^4^, 1 × 10^5^, 1.5 × 10^5^, 9 × 10^5^ or 1 × 10^6^ WT or genetically modified KLN205 and LG1233 tumor cells were subcutaneously injected into the flank of DBA/2J, C57BL/6, TCRα-/-or nude mice. For therapy, tumor-bearing mice were treated daily with 10 mg/kg verubecestat intraperitoneally. To observe the anti-tumor memory response, tumor free mice were rechallenged with 1 × 10^4^ LG1233 cells or 1 × 10^6^ B16F10 cells. To monitor tumor volume, the long and short diameters of tumors were measured with Vernier caliper twice a week after tumors were palpable. Mice were euthanized when tumor volume reached a size of 2,000 mm^3^.

### In vivo depletion and neutralization experiments

The depletion of lymphocytes in mice was conducted by injection of neutralizing antibodies. In brief, C57BL/6 mice were given intraperitoneal injection of antibody against CD4 (100 μg, twice a week, Bio X Cell), CD8 (100 μg, twice a week, Bio X Cell), and NK1.1 (250 μg, once a week, Bio X Cell), respectively. DBA/2J mice were given intraperitoneal injection of antibody against CD4 (100 μg, twice a week, Bio X Cell), CD8 (100 μg, twice a week, Bio X Cell), and asialo-GM1 (50 μg, once a week, Thermo Fisher Scienfic), respectively. All depletion assays were initiated one day before tumor cells inoculation. Specific cell depletion was confirmed by flow cytometry. Neutralization of Cxcr3 was accomplished by intraperitoneal injection of anti-Cxcr3 antibody (200 μg, twice a week, Bio X Cell) one day ahead of tumor injection. Neutralization of Ifnar1 was accomplished by intraperitoneal injection of anti-Ifnar1 antibody (200 μg, twice a week, Bio X Cell) one day ahead of tumor injection. The same amount of isotype IgG were used as the control.

### Immunization

For preventative peptide vaccine, naïve mice were vaccinated intraperitoneally by APP E1-Fc peptide/Aluminium hydroxide gel (Alhydrogel adjuvant 2%, vac-alu-250, InvivoGen), Aβ peptide/CFA, or Aβ peptide/aluminium hydroxide gel, once a week for a total of three weeks. One week after the last immunization, mice were challenged subcutaneously with tumor cells.

### Immunofluorescence

Tumor tissue samples were analyzed for the abundance of intratumoral leukocytes infiltration by immunofluorescence. After fixation with acetone, tumor frozen sections were blocked with serum-free Protein Block (DAKO, X0909) at room temperature for 20 mins. Sections were stained with antibodies against CD45-FITC (1:100, Biolegend), CD8-Alexa Fluor 594 (1:100, Biolegend), CD4-FITC (1:100, Biolegend), or NK1.1-Alexa Fluor 647 (1:100, Biolegend) at 4 °C overnight. Tissues were then counterstained with DAPI for 10 min at room temperature. Sections were mounted with ProLong Gold antifade reagent with DAPI (Invitrogen). Leukocytes were visualized and imaged under fluorescence microscope (Zeiss). The area of lymphocytes in tumor tissues was quantified by ImageJ (FiJi version) software.

### Immunohistochemistry

Human lung adenocarcinoma and squamous carcinoma tissue arrays were obtained from Southwest Hospital (Chongqing, China) and Shanghai Outdo Biotech Company (Shanghai, China). Human breast cancer and colon cancer tissue arrays were obtained from Shanghai Outdo Biotech Company (Shanghai, China). Briefly, formalin-fixed paraffin-embedded tissue sections were deparaffinized and then rehydrated by serial passage through xylene and graded ethanol. Antigen retrieval was performed, followed by blocking buffer incubation at room temperature. Slides were incubated in the primary antibody against APP (1:100, Abcam), BACE-1 (1:100, Abcam), Aβ1-42 (1:100, Abcam) and Aβ1-42 (1:100, Millipore) at 4 °C overnight. After washing out primary antibody, slides were then incubated with secondary antibody-conjugated with HRP at 37 °C, and the immunocomplex was visualized using 3,3’-diaminobenzidine. Then sections were counterstained with hematoxylin. All slides were subsequently imaged using microscope.

### Western blot

Whole cell lysates were prepared with cold RIPA buffer (Sigma-Aldrich) on ice. After quantification, protein sample loading buffer (Invitrogen) was added and the samples were boiled for 5 min at 95 °C. For SDS-PAGE electrophoresis, 20 to 50 μg total protein samples were loaded onto 4%-12% NuPAGE Bis-Tris Gels (Thermo Fisher Scientific). After protein was transferred into PVDF membranes (Millipore-Sigma), membranes were blocked with 5% BSA and then incubated with primary antibody at 4 °C overnight. The following antibodies was used: anti-APP C-terminal (1:1000 Abcam), anti-BACE-1 (1:1000, Abcam), anti-APP KPI (1:1000, Millipore), anti-APP OX2 (1:1000, Millipore), anti-Gsk3β, anti-pIRF3 (S396) (1:1000, Cell Signaling Technology). Anti-GAPDH antibody (1:1000 Santa Cruz) was used as internal control. Blots were then incubated with HRP-conjugated anti-rabbit or anti-mouse secondary antibody (Thermo Fisher Scientific). Signals were detected with SuperSignal West Pico PLUS Chemiluminescent Substrate (Thermo Fisher Scientific).

### qRT-PCR

To induce IFN-I response *in vitro*, cells were treated with 100 ng/ml-10 μg/ml LPS from *Escherichia coli* (serotype O111:B4; Sigma-Aldrich) for 2 hr. Total RNA from cells was extracted and purified with Direct-zol RNA kits (ZYMO Research). RNA was then reversely transcribed into cDNA using iScript cDNA Synthesis Kit (Biorad) or iScript Reverse Transcription Supermix Kit (Quanta Bio). Gene expression was assessed using SYBR Green Supermix (KAPA Biosystems) according to the manufacturers’ protocols. Results were calculated by the 2^-ΛΛCt^ method. Primer sequence was provided in Supplementary Table 1.

### RNA-seq

Total RNA was isolated from LG1233 and KLN205 tumor tissues according to the manufacturer’s instructions (ZYMO Research) and dissolved in RNase-free water. The purity of RNA was evaluated with a NanoPhotometer Spectrophotometer (IMPLEN), while the RNA integrity and quantification were assessed with the RNA Nano 6000 Assay Kit of the Bioanalyzer 2100 system (Agilent Technologies). Library construction and RNA sequencing was performed by the Novogene Corporation. Specifically, libraries were prepared using the NEB Next Ultra RNA Library Prep Kit for Illumina (NEB) according to the manufacturer’s protocol. Libraries were sequenced using an Illumina HiSeq machine and paired-end reads were generated.

### Flow cytometry

Tumor tissues were firstly disaggregated mechanically and digested with 1 mg/ml collagenase I (GIBCO) at 37 °C for 1 hr. Single cell suspension was then subjected to red blood cell lysis using ACK lysing buffer, followed by washing with PBS. Cells were first stained with violet-or NearIR-LIVE/DEAD fixable dead cell stain dye (1:2000, Invitrogen), followed by incubation with the following antibodies for 30 min at 4 °C: CD45–FITC, CD4–FITC, CD4 –PE–Cy7, CD8 –PE–Cy7, CD8-PerCP-Cy5.5, TCRβ–PE, NK1.1–APC, Ly6G–APC, CD11b–PerCP–Cy5.5, CD11c–PE–Cy5 and F4/80–PE (Biolegend, 1:100). For tumor cell phospho-flow cytometry, LG1233 tumor cells were fixed and permeabilized in True-Phos Perm Buffer (Biolegend) for 1 hr at –20°C. Cells were then stained with p-TBK1 (S172)-PE (Cell Signaling Technology, 1:100), p-IRF3 (S396)-PE (Cell Signaling Technology, 1:100), or p-Stat1-Alexa Fluor 488 (Y701) (BD Bioscience, 1:100) for 30 min at room temperature. For intracellular staining, tumor cells were fixed and permeabilized, followed by staining with anti-human IgG Fc-PE-Cy7 (Biolegend, 1:100) for 30 min at room temperature. Samples were acquired on a BD FACSCanto II machine (BD Biosciences) and data were analyzed using FlowJo software.

### Bioinformatics analyses

For bulk RNA-seq analysis, raw short reads were filtered and cleaned by *fastp* v0.20.1 in paired-end mode^60^. Sequence adapters and low-quality reads were trimmed with ‘-q 20’ parameter. The resulting clean reads were aligned to the Gencode GRCm38 reference genome (release v25) using STAR v2.7.5a^61^ with default parameters. Aligned reads with a mapping quality score >20 were retained using Samtools v1.18^62^ with the parameters “-f 2 –q 20”. The number of mapped fragments per gene for each sample was quantified using *featureCounts* v2.0.1^63^. Differential gene expression (DEG) analysis was conducted using *DESeq2* v1.36.0^64^. *DESeq2* uses a negative binomial distribution and a generalized linear model to model gene expression data. Genes with >5 reads in at least 3 samples were retained during the analysis. Gene count normalization was performed using variance stabilizing transformation (VST) method implemented in *DESeq2*. P values were assessed using Wald test and then adjusted for multiple testing corrections using the Benjamini-Hochberg method. DEGs with an adjusted p value < 0.05 were selected for subsequent analysis. Hierarchical clustering and heatmaps were constructed using R *pheatmap* v1.0.12, which uses the Euclidean distance method to assess sample and gene correlation. To visualize DEGs, volcano plots were generated using the R *EnhancedVolcano* v1.14.0 package. Enrichment of functional pathway analysis of significant DEGs was performed using R package *clusterProfiler* v4.4.4^65^. The gene set enrichment analysis (GSEA) was conducted using *fgsea* software. To assess the difference of immune response between APP-proficient and APP-deficient tumor samples, we performed immune deconvolution analysis with the R package *GSVA*^66^ using ssGSEA method with default parameters. The gene markers of each immune pathway and cell type were retrieved from a previous study^67^. *GEPIA2*^28^ was used to evaluate the correlation between APP gene expression and patient clinical outcome.

### Statistical methods

Statistical analysis was performed using GraphPad Prism 8 software (GraphPad Software). Statistical analysis for bioinformatics was performed in R v4.0.3. For survival benefits assessment, Kaplan-Meier methods and log-rank test were used to calculate statistical significance. Wilcox test or two-tailed t-tests were used to compare statistical difference between two groups. ANOVA models were used to compare outcomes across multiple experimental groups. For multiple comparisons, Tukey corrections were used to adjust P values. Statistical significance was considered as *P* < 0.05.

## Figure legends

**Extended Data Fig. 1.** APP and its β-cleavage products in cancer tissues. A, Immunohistochemical staining of APP in human lung cancers. Quantification of APP expression in tumor and stroma area (n=30 for LUSC, n=26 for LUAD). B-C, Immunohistochemical staining of Aβ1-42 in human LUSC (B) and LUAD (C) using antibody from Millipore and Abcam company. D-E, Immunohistochemical staining of Aβ1-42 in a second cohort of patients with human LUSC (D) and LUAD (E) (n=90 for LUSC, n=92 for LUAD). F-G, Immunohistochemical staining of Aβ1-42 in patients with colon cancer (F) (n=100) and breast cancer (G) (n=143). Scale bars, 50 μm. H, Quantification of Aβ1-42 expression in different types of cancer tissues as in (D-G). I-J, The expression of APP and BACE-1 was assessed in human (I) and murine (J) lung cancer cells. K, Schematics of protein domain of APP 770. Data are mean ± s.d. \*\*\*\**P* < 0.0001.

**Extended Data Fig. 2.** APP promotes tumor growth. A-B, KLN205 cells (A) were injected subcutaneously into DBA/2J mice. LG1233 cells (B) were inoculated subcutaneously into C57BL/6J mice. All mice were treated daily with verubecestat intraperitoneally. Tumor growth curve was plotted. C-D, DBA/2J mice were prophylactically vaccinated with Aβ1-42 and adjuvant CFA (C) or Aβ1-5/Aβ1-16 along with adjuvant alum (D), once a week for 3 weeks. Mice were then challenged with KLN205 cells. Tumor volume was tracked. E-H, The relationship of Aβ1-42 expression and prognosis in indicated cancer patients. I-L, KLN205 (I) and LG1233 cells (K) were transfected with none targeting shRNA (shNT) and shRNA targeting APP (shAPP). Knockdown efficiency was evaluated with Western blot. The in vitro proliferation curve of shNT and shAPP KLN205 (J) and LG1233 cells (L). M-N, shNT and shAPP KLN205 (M) and LG1233 cells (N) were injected subcutaneously into DBA/2J and C57BL/6J mice, respectively. Tumor volume was measured. O-P, CRISPR-Cas 9 indel events near target site was confirmed by sequencing PCR products. Q-R, the expression of APP’s domain-Fc in KLN205 APP sgRNA-1 (Q) and LG1233 APP sgRNA-1 (R) cells was evaluated by flow cytometry. Data are mean ± s.d. \*\**P* < 0.01.

**Extended Data Fig. 3.** APP-deficient tumor features and APP excludes lymphocytes in lung cancer. A-B, LG1233 cells with control or APP sgRNA-1 were injected into male C57BL/6J mice. KLN205 cells with shNT or shAPP were injected into male DBA/2J mice. Bulk tumor RNA-seq was performed. GSEA analysis of LG1233 (A) or KLN205 (B) tumor tissues. C-E, The percentage of CD45+ cells were analyzed by flow cytometry in Ctrl or APP sgRNA-1 LG1233 (C, D) and KLN205 (E) tumor tissues. F, Control or APP sgRNA-1 LG1233 cells were subcutaneously injected into C57BL/6J mice. The percentage of lymphocytes in TME were analyzed by flow cytometry. n=7. G-H, The immune deconvolution of lung cancer patients with high and low expression of *APP* in TCGA data portal cohort. Patients were divided into high and low-expression groups according to a quartile cut-off value. Data are presented as means ± s.d. \**P* < 0.05, \*\**P* < 0.01, \*\*\**P* < 0.001, \*\*\*\**P* < 0.0001.

**Extended Data Fig. 4.** Lymphocyte depletion experiments in lung cancer models and APP deprivation induces anti-tumor memory response. A-C, Control or APP sgRNA-1 LG1233 cells were subcutaneously injected into C57BL/6J mice. Lymphocyte depletion efficiencies by anti-CD4, anti-CD8 and anti-NK1.1 antibodies in LG1233-bearing mice were assessed by flow cytometry. Representative plots (A, B) and statistics (C) are shown. D, Control or APP sgRNA-1 KLN205 cells were subcutaneously injected into DBA/2J mice. Lymphocyte depletion efficiencies by anti-CD4, anti-CD8 and anti-asialo-GM1 antibodies in KLN205-bearing mice were assessed by flow cytometry. E-F, APP sgRNA-1 LG1233 tumor-free mice were rechallenged with parental LG1233 tumor cells. Naïve C57BL/6 mice as control. Tumor volume was measured and survival curve was plotted. n=5. G, LG1233 tumor-free C57BL/6 mice were rechallenged with B16 tumor cells. Tumor volume was measured. n=5. \*\**P* < 0.01, \*\*\**P* < 0.001, \*\*\*\**P* < 0.0001.

**Extended Data Fig. 5.** APP’ E1 domain suppresses type I interferon response. A, The expression of phosphorylated IRF3 was assessed by western blot in LG1233 cells. B, KLN205 WT tumor cells were treated with recombinant APP E1-Fc peptide for 12 hr, followed by treatment with LPS for 2 hr. The expression of IFN-I was assessed by RT-PCR. C, KLN205 WT tumor cells were treated with recombinant Fc peptide for 12 hr, followed by treatment with LPS for 2 hr. The expression of IFN-I was assessed by RT-PCR. D, LG1233 WT tumor cells with mock-GFP or APP E1-Fc-GFP overexpression were treated with LPS for 2 hr. The expression of IFN-I was assessed by RT-PCR. E, KLN205 WT tumor cells with mock-GFP or APP E1-Fc-GFP overexpression were treated with LPS for 2 hr. The expression of IFN-I was assessed by RT-PCR. F-G, Human LUSC H520 cells (F) and human LUAD HCC827 cells (G) were treated with recombinant APP E1-Fc peptide for 12 hr, followed by treatment with LPS for 2 hr. The expression of IFN-I was assessed by RT-PCR. H, LG1233 WT tumor cells were treated with recombinant sAPPα or sAPPβ peptide for 12 hr, followed by treatment with LPS for 2 hr. The expression of IFN-I and related genes was evaluated by RT-PCR. Data are presented as means ± s.d. \**P* < 0.05, \*\**P* < 0.01, \*\*\**P* < 0.001, \*\*\*\**P* < 0.0001.

**Extended Data Fig. 6.** Gsk3β mediates the suppressive effect of APP’ E1 domain on type I interferon response. A-B, The expression of GFP in KLN205 (A) and LG1233 (B) tumor cells after transfection with indicated MSCV-vectors. Co-expression of GFP was used as a selection marker in stable cell lines. C, The expression of IFN-I in LG1233 tumor cells with mock-GFP or Gsk3β S9A-GFP overexpression was assessed by RT-PCR in the absence of LPS. D, The expression level of p-TBK1 (S172) in Gsk3β S9A overexpressed LG1233 cells, as assessed by flow cytometry. E, GO pathway analysis of KLN205 tumors with Gsk3β S9A overexpression. n=3. F-G, Gene set enrichment analysis of RNA-seq data from bulk KLN205 tumor tissue with Gsk3β S9A overexpression. n=3. H, The growth curve of KLN205 tumors with Gsk3β S9A overexpression. I-J, Knockdown efficiency of shGsk3β in APP sgRNA-1 LG1233 (I) and APP sgRNA-1 KLN205 (J) cells. K-M, KLN205 tumor cells infected with MSCV-mock-GFP (K), MSCV-Gsk3β S9A-GFP (L) and MSCV-Gsk3β K85A-GFP (M) vectors, were treated with recombinant APP E1-Fc peptide for 12 hr, followed by treatment with LPS for 2 hr. The expression of IFN-I and related genes was evaluated by RT-PCR. Data are presented as means ± s.d. \**P* < 0.05, \*\**P* < 0.01, \*\*\**P* < 0.001, \*\*\*\**P* < 0.0001.

