## Supplementary figures and images for "Amyloid-β precursor protein promotes tumor growth by establishing an immune-exclusive tumor microenvironment"

### Supplementary Figure 1

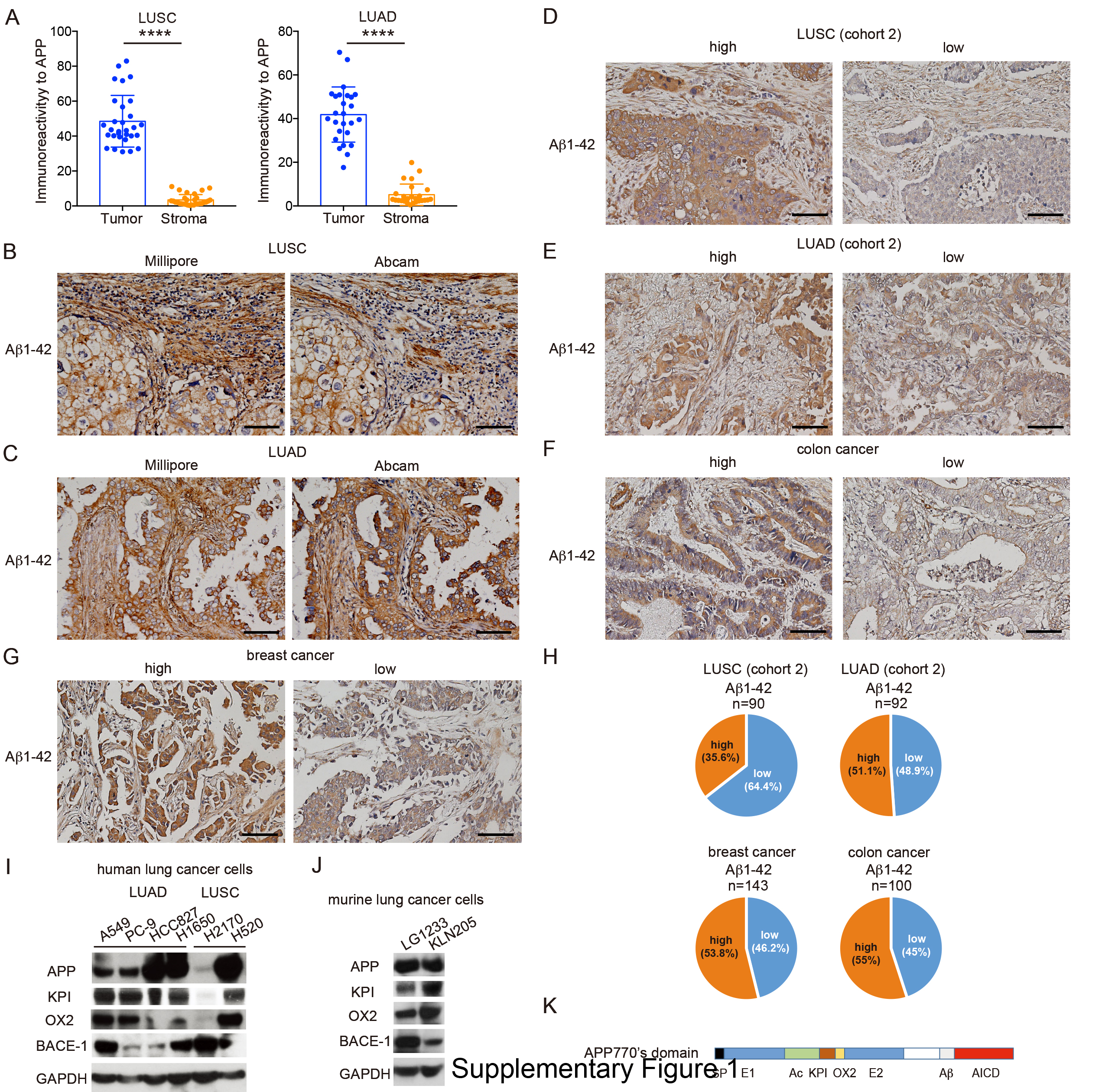

### Supplementary Figure 2

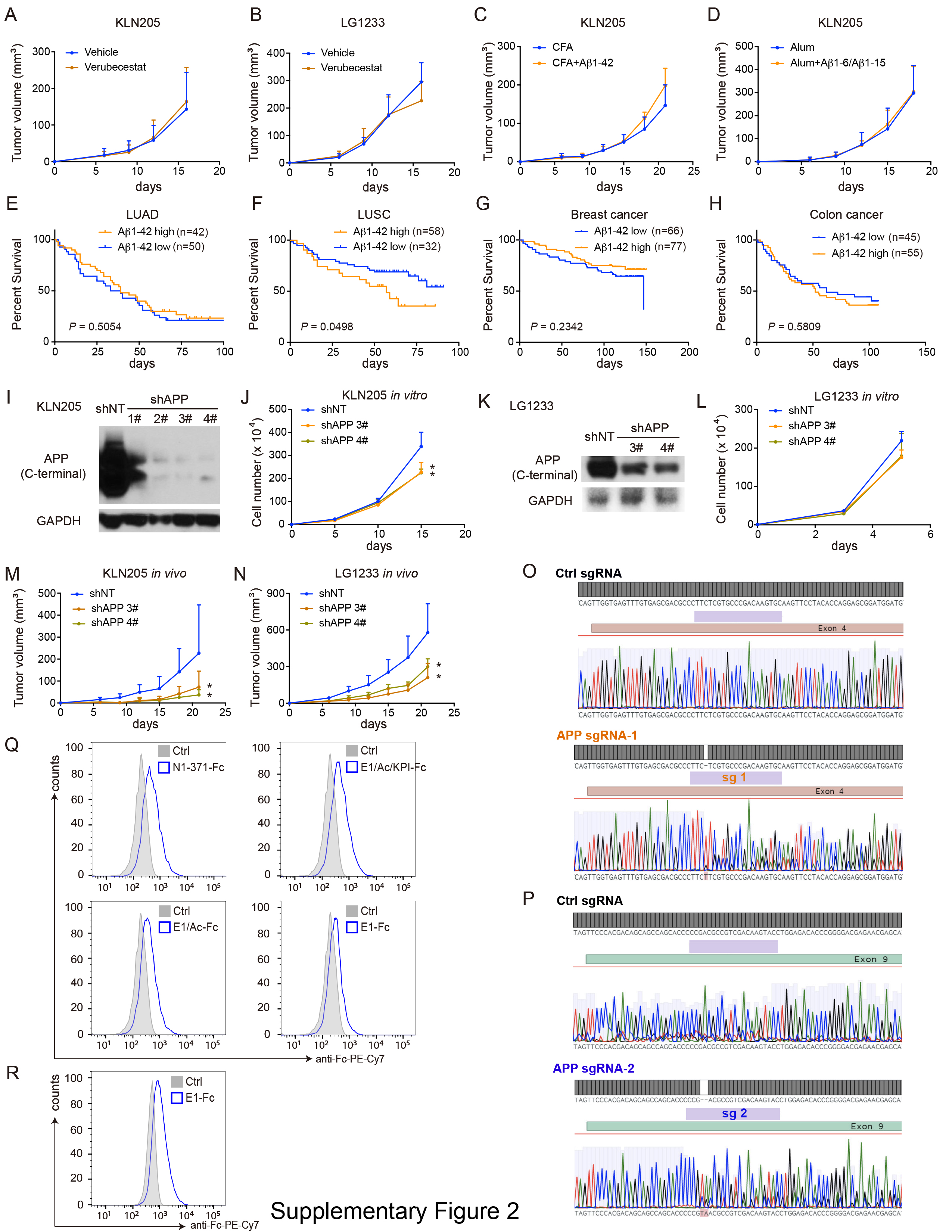

### Supplementary Figure 3

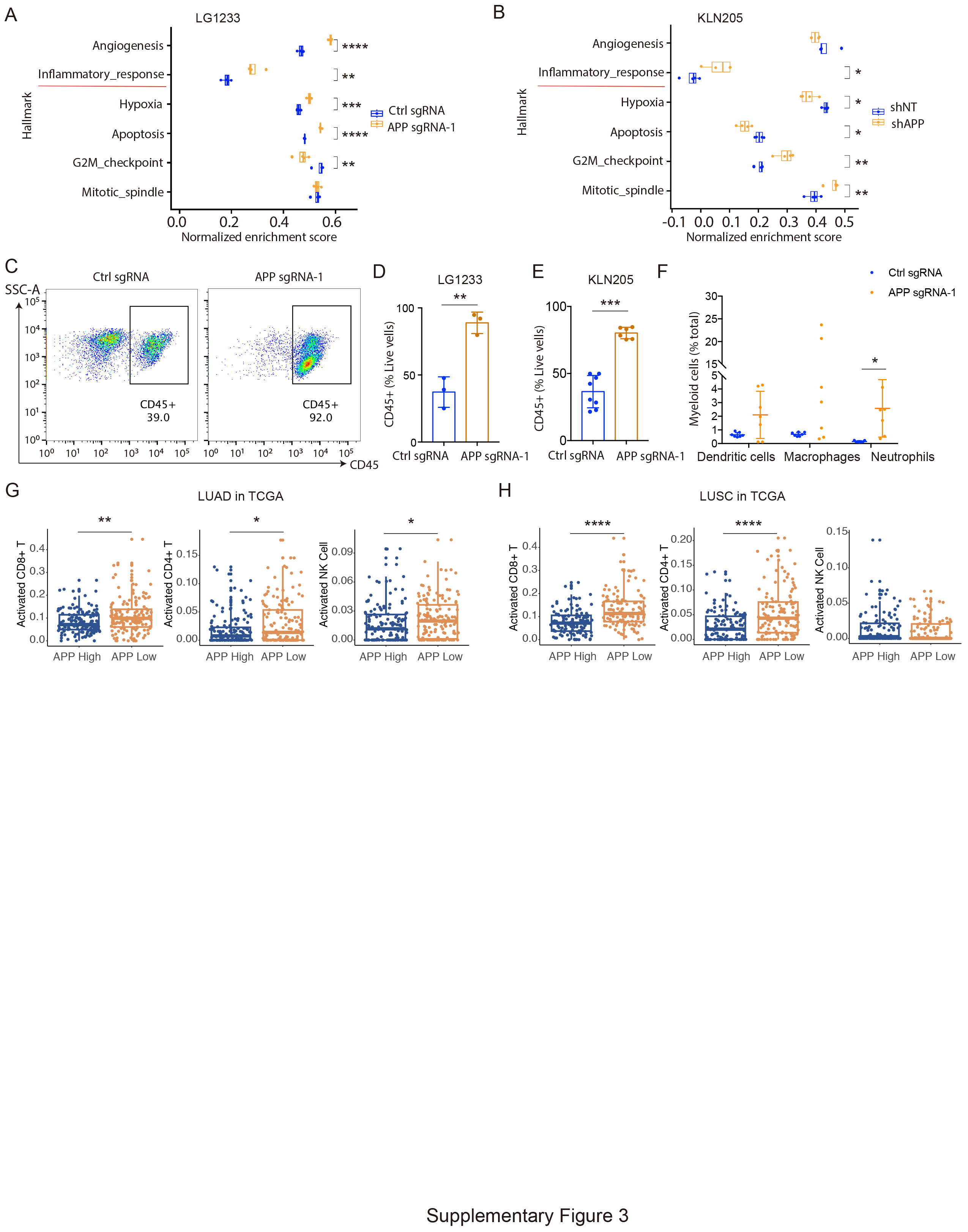

### Supplementary Figure 4

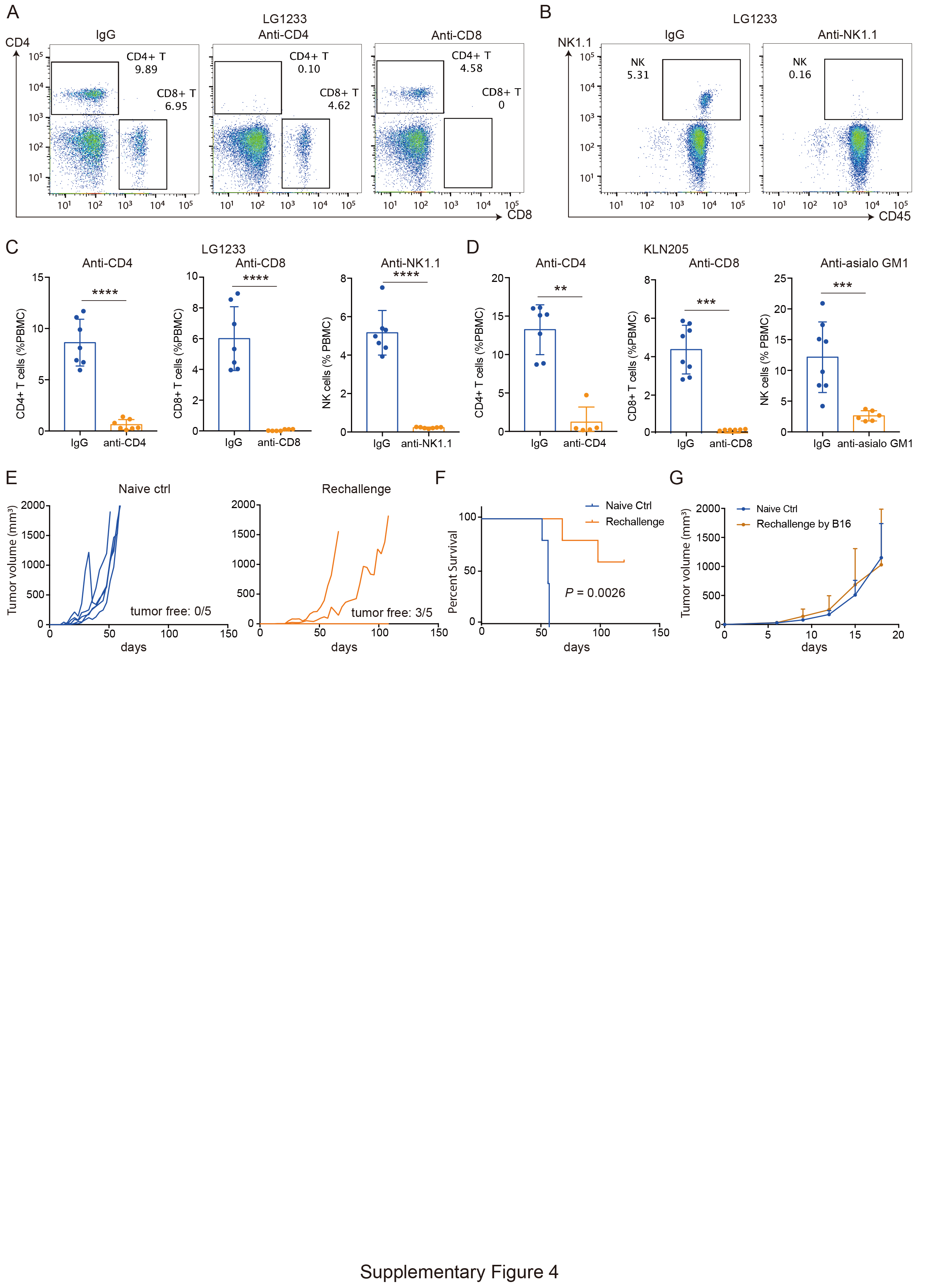

### Supplementary Figure 5

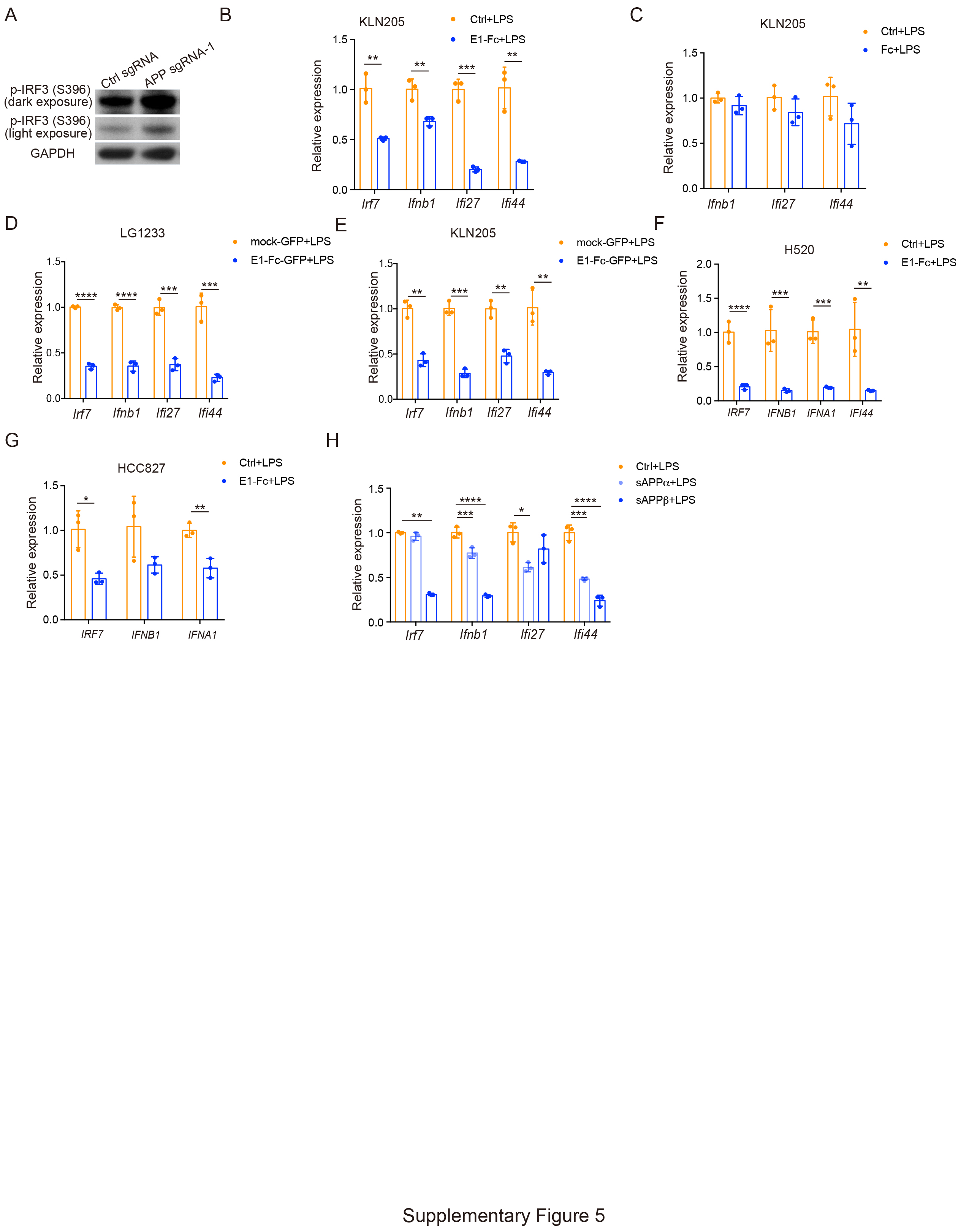

### Supplementary Figure 6

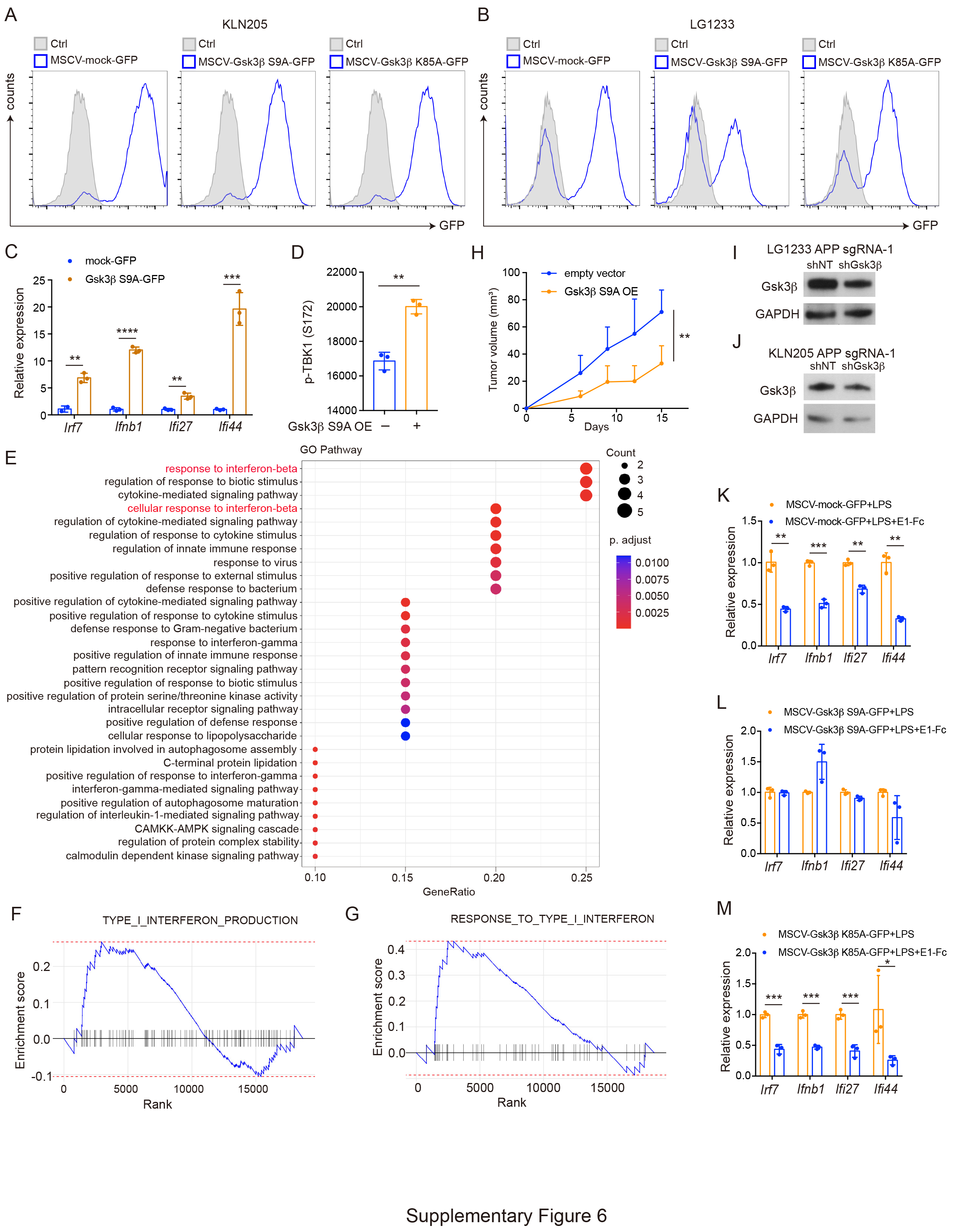
